# Comparative genomics reveals phylogenetic intermixing of *Stomoxys* fly, manure, and bovine mastitis-associated bacteria in dairy settings

**DOI:** 10.1101/2025.01.31.635862

**Authors:** Andrew J. Sommer, Travis K. Worley, Panagiotis Sapountzis, Kerri L. Coon

## Abstract

*Stomoxys* flies are highly ubiquitous blood-feeding pests on dairy farms and are suspected carriers of pathogenic bacteria due to their close association with both manure reservoirs and cattle hosts. While prior studies using amplicon sequencing and culture dependent methodologies have been used to characterize the composition of the *Stomoxys* microbiota, little is known about strain level genomic and functional diversity of *Stomoxys*-associated bacteria. In this study, we address this key gap in knowledge by using whole genome sequencing to provide the first comparative genomic analysis of *Stomoxys-*derived *Escherichia coli*, *Klebsiella pneumoniae*, and *Staphylococcaceae* isolates. Our results show that fly and manure isolates collected from the same farm system are phylogenetically interspersed, with subsequent pairwise genome alignments revealing near identical strains and plasmids shared between the two sources. We further identified a phylogenetic clade of *Mammaliicoccus sciuri*, which contains known mastitis agents that were associated with both flies and manure. Functional analysis revealed that this clade was highly enriched in xylose metabolism genes that were rare across other *M. sciuri* lineages, suggesting potential niche differentiation within the genus. Collectively, our results provide strong evidence for the acquisition of fecal-associated bacterial isolates by adult *Stomoxys* flies, confirming the link between biting muscid flies and manure habitats. The intermixing of fly and manure isolates in clinically relevant taxonomic groups strongly suggests that flies serve as carriers of opportunistic mastitis-causing or other fecal-borne pathogens, and that flies may serve as important vehicles of pathogen dissemination across the dairy farm environment.

**Author summary:** Blood-feeding *Stomoxys* flies are a major pest species in dairy operations and have been implicated as carriers of fecal-borne pathogens responsible for bovine mastitis and other important cattle diseases. In a previous study, we generated a large collection of bacterial isolates from manure and *Stomoxys* flies collected from dairy barns. Most bacterial isolates derived from flies were identified as taxa associated with bovine mastitis, including *Escherichia coli*, *Klebsiella pneumoniae*, and non-aureus *Staphylococcus*; however, it is unknown if these represent bacterial strains acquired by flies from manure. Here, we examined this possibility directly by performing whole genome sequencing of a subset of potential mastitis pathogens derived primarily from flies and manure and supplemented this analysis with publicly available genomes on NCBI (including mastitis and additional fly and manure strains). Our results reveal significant interrelations between fly, manure, and clinically derived mastitis strains, suggesting that *Stomoxys* flies participate in the carriage and dissemination of bovine mastitis pathogens acquired from manure. Additionally, this work highlights the functional diversity of bacterial pathogens within the barn environment and greatly expands the available genomic data for understudied mastitis pathogens such as *Mammaliicoccus sciuri*.

## Introduction

The effective control of infectious diseases across agricultural systems is crucial for ensuring the economic stability of the global food system. On commercial dairy farms, strict hygiene practices and bio-control programs are often implemented to limit the spread of animal-borne pathogens to farm workers, other livestock, or through the food system to consumers (1–3). While much attention has been given to the surveillance of pathogens in cattle or food products (4–6), far less is known about the role of pest species, including insects, rodents, and birds, in the maintenance or potential transmission of microorganisms through the farm environment.

*Stomoxys* flies are ubiquitous blood-feeding pests inhabiting dairy farms across the globe and have been previously implicated as carriers of pathogenic bacteria on dairy farms due to their close association with both bovine manure and cattle hosts (7–12). Adult female flies preferentially oviposit eggs into aged cattle manure, which serves as a protein, carbohydrate, and microbe rich source of nutrition for developing larvae (13–16). As adults, both male and female flies require daily nutritional bloodmeals from cattle or other mammalian hosts (12,17). Ingested bacteria can be excreted by *Stomoxys* flies via regurgitation during blood-meals (18,19), or further dispersed through the environment by external carriage or defecation post-feeding (11,20). Defensive behaviors by livestock during feeding will often lead to multiple interrupted blood-meals (17,21,22), facilitating further contact between *Stomoxys* flies and multiple bovine hosts. Recently, we performed the first culture independent (16S rRNA gene amplicon sequencing) characterization of bacterial communities associated with adult *Stomoxys* flies and bovine manure samples collected longitudinally in a working dairy facility. Our results showed that the *Stomoxys* microbiota is primarily comprised of *Enterobacteriaceae*, *Staphylococcaceae*, and other opportunistic pathogenic bacterial taxa (7), which can be readily isolated from both the exogenous and endogenous surfaces of adult flies (8). These bacteria are the most common causative agents of environmental bovine mastitis (opportunistic intramammary infections) (1,23), which can be spread through exposure to manure or contaminated bedding. We identified that the majority of bacterial phylotypes sequenced from flies were also found in low relative abundance in manure samples, suggesting the potential for the acquisition of manure and mastitis-associated bacteria by biting flies on dairy farms.

In this study, we performed short-read genome sequencing of a collection of bacterial strains derived primarily from flies and manure samples collected from a dairy facility in Wisconsin, USA. Comparative genomic analysis was performed on 298 sequenced bacterial genomes across four focal taxa (*Escherichia coli*, *Klebsiella pneumoniae*, *Mammaliicoccus sciuri,* and *Staphylococcus spp.*) associated with animal (bovine mastitis) and zoonotic diseases. We then constructed larger reference genome sets to compare the phylogenetic and functional diversity of fly-, manure-, and mastitis-derived strains with bacteria isolated from other hosts and environments.

Our results reveal phylogenetic intermixing between fly- and manure-derived isolates as well as the presence of near-identical strains across sample types, suggesting a role in the dispersal of manure bacteria and mobile genetic elements by flies on dairy farms. Subsequent analysis of *M. sciuri* indicated that fly-, manure-, and mastitis-derived strains were strongly associated with a specific clade of *M. sciuri*, which was enriched in xylose metabolism genes (*xylA*, *xylB*) that were absent from other *M. sciuri* lineages. Collectively, these results provide strong evidence for the interconnectedness of fly and manure microbiomes and further suggest a role for flies in the carriage of mastitis pathogens in dairy settings.

## Results

### Development of a bacterial isolate collection

In three previous studies, we have worked to develop a bacterial isolate collection of fly- and manure-derived bacterial strains isolated from samples collected across two focal dairy farms in south central Wisconsin (7,8,24). This collection contains about 1200 bacterial isolates, of which 80 manure-derived and 208 fly-derived isolates were processed in this study for short-read whole genome sequencing. We additionally sequenced 6 *K. pneumoniae*, 1 *E. coli*, and 1 *S. haemolyticus* isolates, which represented known clinical mastitis isolates collected by the Wisconsin Veterinary Diagnostic Laboratory (WVDL) from mastitic cows housed in the same facilities.

### Collection of draft genomes from Stomoxys flies and the barn environment

A total of 296 genomes, including 208 fly-, 80 manure-, and 8 mastitis-derived isolates were sequenced as part of this study (Table S1). Draft genome assembly statistics were determined via Quast (25), with a median N50 metric of 285,278 base pairs (bp) across all sequenced isolates (Figure 1A). We used the BUSCO (Benchmarking Universal Single-Copy Orthologs) to search for universal single copy orthologs across assembled genomes and found that 221 draft isolates contained all 124 expected orthologs (26). A range of 118 to 123 universal single copies were found across the remaining assemblies (Figure 1B). Genome completeness and contamination were calculated using CheckM with a full reference tree, which showed a median completeness of 99.45% (95.91% to 100%) and a median contamination level of 0.735% (0% to 4.38%) (Figure 1C-D). Taxonomic assignment was performed using PubMLST Ribosomal Multilocus Sequence Typing (rMLST), which revealed that the most common bacterial taxa in the collection included strains of *Escherichia coli, Klebsiella pneumoniae, Staphylococcus*, and *Mammaliicoccus* (Table S1; Figure 1E). All assembled genomes in this study were then screened for the presence of Antimicrobial Resistance Genes (ARGs), Virulence Factors (VFs), and plasmid replicon genes (Table S2), as appropriate (27–31).

**Figure 1.**
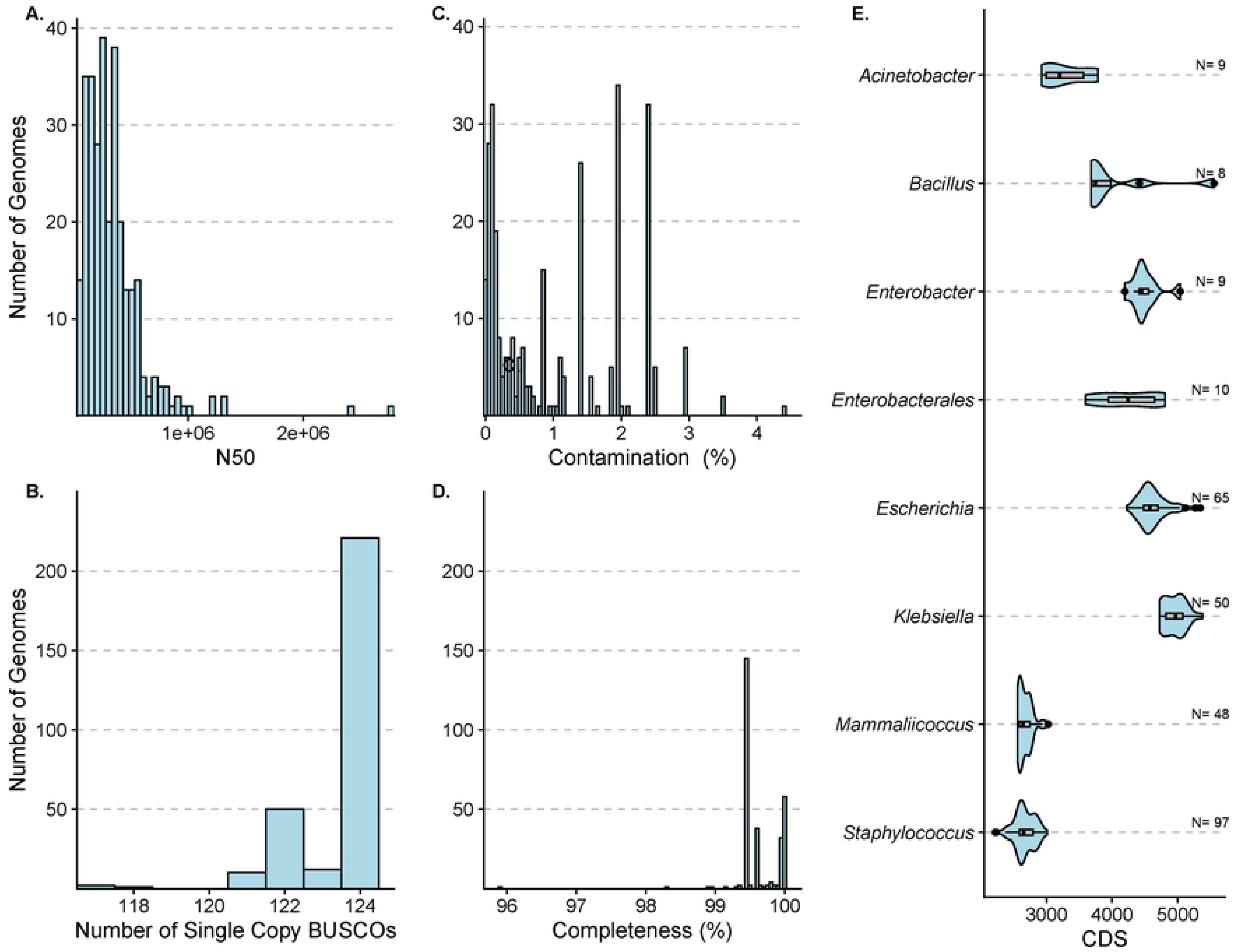
Genome assembly statistics. (A) Histogram depicting the distribution of N50 contig lengths across sequenced genomes. (B) Histogram depicting the number of BUSCO (Benchmarking Universal Single-Copy Orthologs) detected across sequenced genomes. (C/D) Histogram depicting the calculated contamination (C) and completeness (D) levels across sequenced genomes. (E) Violin plots showing the distribution of CDS (coding sequences) across sequenced taxonomic groups. N represents the number of genomes within each taxonomic group.

### Analysis of sequenced E. coli isolates

We performed a phylogenetic analysis of the 65 *E. coli* isolates sequenced in this study, which included 36 *Stomoxys*-derived *E. coli,* 28 manure-derived *E. coli* isolates, and 1 mastitis-derived *E. coli* isolate. A maximum-likelihood phylogenetic tree was constructed based on a core genome alignment to show the local population structure of sequenced *E. coli* isolates (Figure 2A). Bacterial isolates clustered based on calculated MASH phylogroups (32), with the majority of isolates belonging to phylogroup B1 (Figure 2A). Conversely, only a single isolate (manure-derived) in the collection belonged to phylogroup B2, and a small minority of strains belonged to phylogroup D. We observed phylogenetic intermixing between fly- and manure-derived *E. coli* isolates on the phylogeny, which included two sets of fly and manure isolate pairs with the same sequence type (ST). The single mastitis-derived *E. coli* isolate also shared the same sequence type (ST2) as the fly and manure isolates, although all three strains had different predicted serotypes (Table S3).

**Figure 2.**
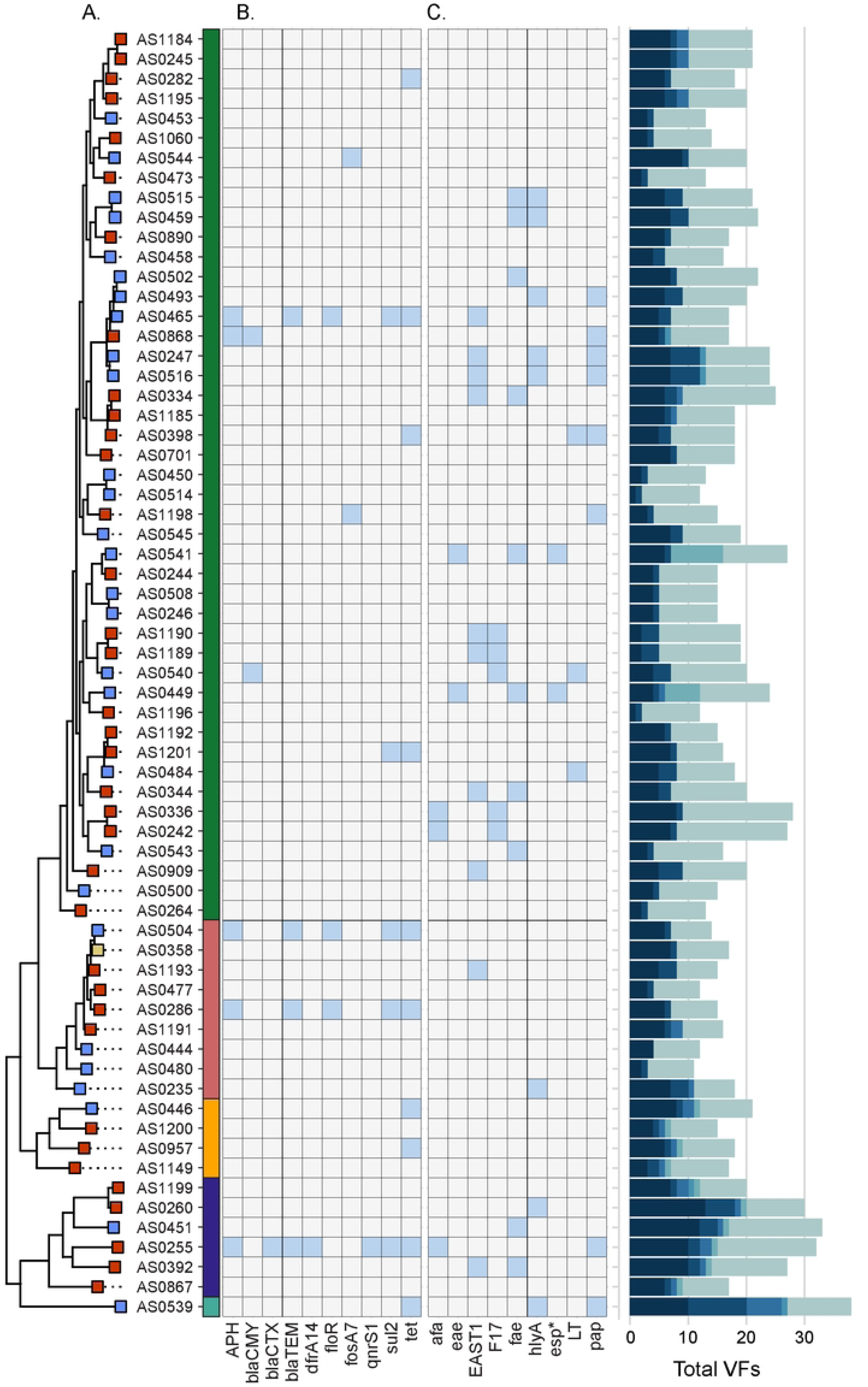
Phylogenetic intermixing of sequenced *E. coli* strains. (A) Midpoint rooted maximum-likelihood phylogenetic tree based on core gene alignment of sequenced *E. coli* isolates. The rectangular node left of the strain name indicates the origin of each isolate as follows: flies (red), manure (blue), mastitis (yellow). The colored bar to the right of the fly strain indicates the MASH phylogroup of each: D (Dark blue; top), B2 (cyan), E (orange), A (light red), B1 (green, bottom). (B) Heatmap depicting the presence-absence matrix (blue indicates presence) of selected ARGs (Resfinder) and VFs (VirulenceFinder). * *esp* refers to strains positive for either *espA*, *espB*, or *espJ*. (C) Bar graph showing the total number of VFs detected in each strain. Stacked colors represent counts of different classes of VFs as follows from light to dark: adhesion, effector delivery system (EFD), invasion, iron uptake, toxin, other VFs.

*E. coli* draft assemblies were screened for the presence of acquired ARGs and VFs using the ResFinder and VirulenceFinder databases, respectively (30,31). Acquired ARG carriage was overall rare throughout the collection; however, we did find both fly- and manure-derived *E. coli* strains that harbored extended spectrum beta-lactamase genes (ESBLs), including *bla*-CTX, *bla*-CMY, and *bla*-TEM (Figure 2B). Carriage of ESBLs was not confined to a specific phylogroup. Virulence factor carriage was mixed throughout the phylogenetic tree, with fly and manure strains carrying a variety of adhesins, toxins, and iron uptake genes (Figure 2C). This notably included four fly-derived and one manure-derived *E. coli* strains encoding genes related to yersiniabactin siderophore production (*irp*, *fyuA*, *ybt*) (Table S2; Table S3). None of the sequenced strains from either manure or flies carried the Shiga (*stx*) toxin, but two phylogroup B1 manure-derived strains encoded *esp* genes (*espA*, *espB*, *espJ*), which are associated with an enterocyte effacement (LEE4) pathogenicity island and several other strains also carried the T3SS effector-like protein gene *espY2* (Table S3) (33,34). Adhesive fimbriae genes found in the sequenced strains included the F17 fimbriae, *pap* fimbriae, and Afa/Dr type (*afa*) adhesins.

We then constructed a larger reference phylogeny to determine the relationship of the sequenced *E. coli* to the global *E. coli* phylogeny. This maximum-likelihood phylogenetic tree was based on a core genome alignment of 410 *E. coli* genomes (Figure 3), which notably included 126 publicly available fly-derived and 121 publicly available mastitis-derived genomes. NCBI records indicate that the fly-derived bacterial strains were mainly isolated from *M. domestica* (Family: Muscidae) or from an unidentified fly; a subset of bacterial strains were identified as being isolated from *Chrysomya* (Family: Calliphoridae) (Table S4). These fly-derived strains placed along the entire *E. coli* phylogeny, with the largest number of strains belonging to phylogroup A (n= 89) followed by B1 (n= 33). A similar pattern was observed for bovine mastitis strains with 84 genomes identified as MASH phylogroup A and 28 genomes identified as phylogroup B1. Pangenome analysis revealed clustering of strains by phylogroup based on gene-content (Figure S1).

**Figure 3.**
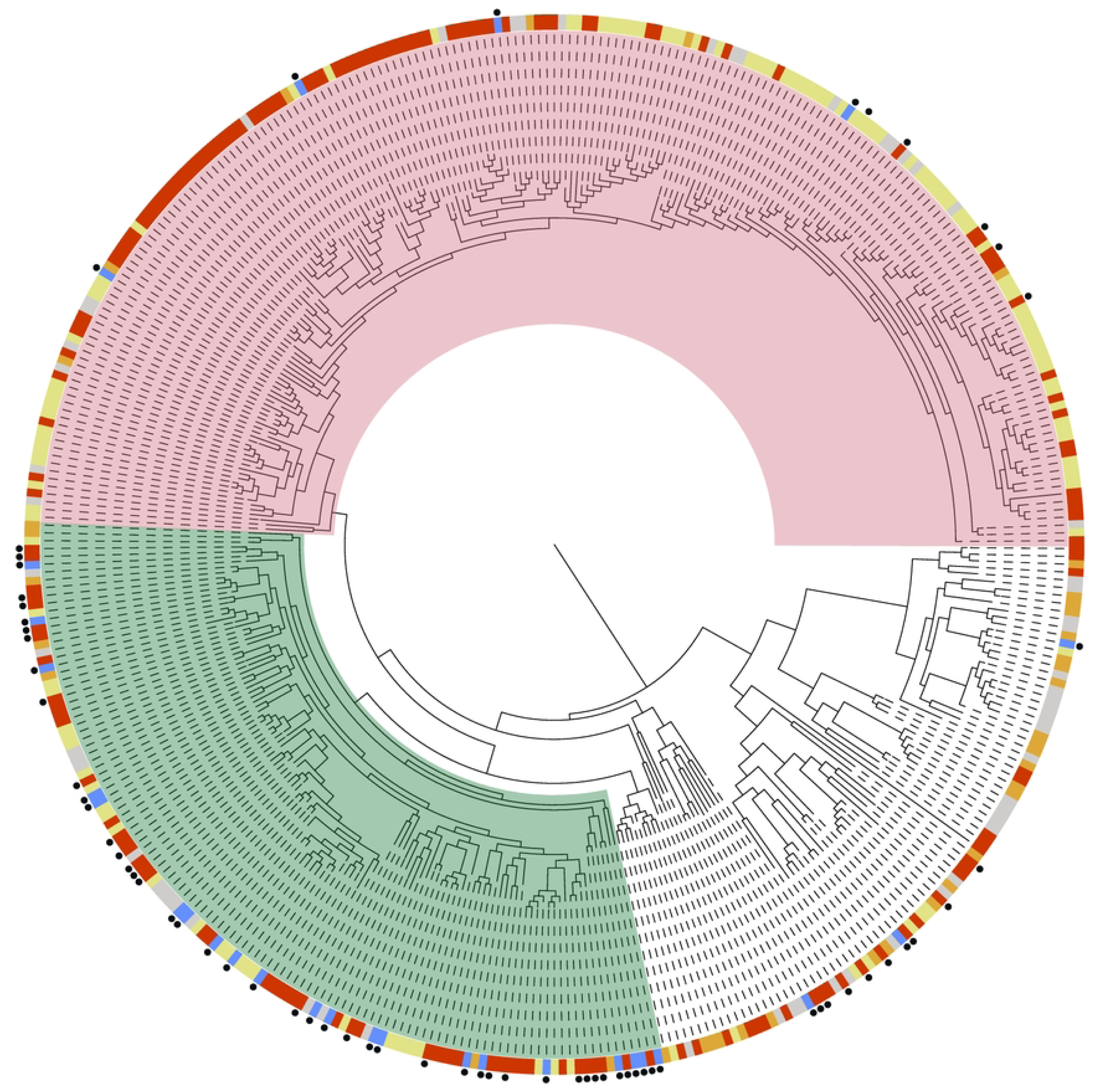
*E. coli* population structure and phylogenic placement of sequenced *E. coli* strains. A midpoint rooted maximum-likelihood phylogenetic tree was constructed based on a core gene alignment of 410 *E. coli* genomes. Isolates sequenced in this study are marked with a black circle. The node color indicates the origin of each isolate as follows: flies (red), manure (blue), mastitis (yellow), other host (nonclinical) (grey), other host (clinical) (orange). The phylogroup A *E. coli* clade is highlighted in red; the phylogroup B1 *E. coli* clade is highlighted in green.

### Analysis of sequenced K. pneumoniae isolates

We constructed a maximum-likelihood phylogenetic tree based on a core genome alignment of 46 sequenced *K. pneumoniae* isolates (21 fly-derived, 19 manure-derived, 6 mastitis-derived), which showed that these isolates formed three distinct lineages with sample origins intermixed within each lineage (Figure 4A). Carriage of acquired ARGs was largely rare, but notably included two fly-derived strains and two manure-derived strains carrying ESBLs (Figure 4B). In contrast to *E. coli*, there was little variation in VFs by *K. pneumoniae* strains and no strains encoded gene clusters associated with hypervirulence (*ybt*, *clb iuc*, *iro*, *rmp*) (Figure 4C; Table S5) (35). *In silico* capsular polysaccharide (CPS) of sequenced *K. pneumoniae* indicated the presence of 33 K-locus types, but only KL14 and KL151 were detected in both a fly and a manure strain (Figure 4C). In both cases, differing sequence types (ST) were identified in the corresponding fly and manure strains (Table S5). KL106 was detected in both a manure and a mastitis-derived strain, which clustered together in the phylogeny and shared the same ST (ST101).

**Figure 4.**
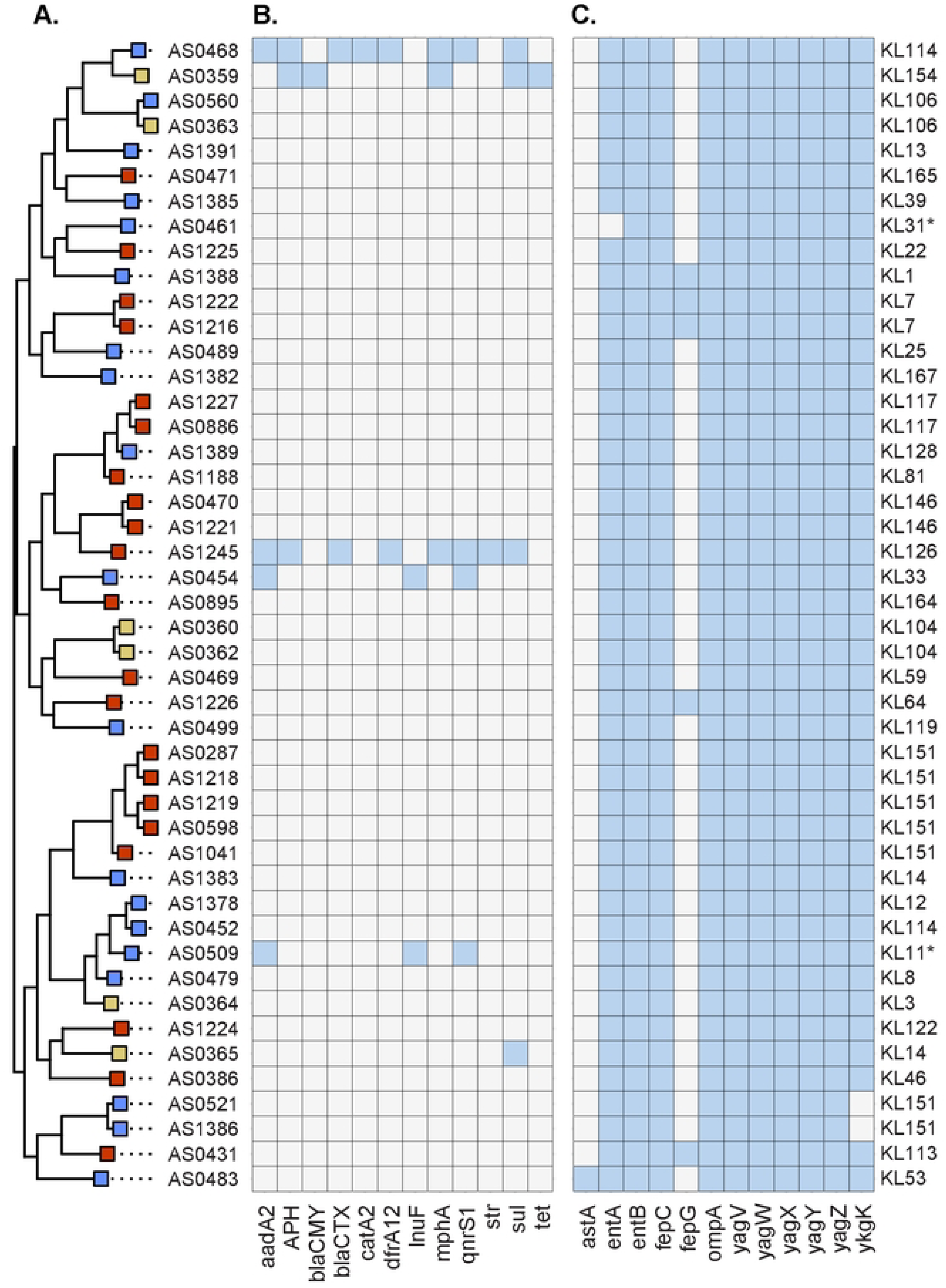
Phylogenetic intermixing of sequenced *K. pneumoniae* strains. (A) Midpoint rooted maximum-likelihood phylogenetic tree based on core gene alignment of sequenced *K. pneumoniae* isolates. The rectangular node left of the strain name indicates the origin of each isolate as follows: flies (red), manure (blue), mastitis (yellow). (B/C) Heatmap depicting the presence-absence matrix (blue indicates presence) of selected ARGs (Resfinder) (B) and VFs (VFDB) (C). The predicated capsular polysaccharide type is provided to the right of the heatmaps.

We then built a reference genome set of 281 *K. pneumoniae* strains, which contained 80 mastitis-derived and 13 fly-derived strains (Table S4). The resulting phylogenetic tree confirmed that flies harbored diverse lineages of *K. pneumoniae* (Figure 5). Despite the presence of deep branching *K. pneumoniae* lineages, pangenome analysis of the reference set revealed no distinct clustering of isolates or lineages by gene content (Figure S2).

**Figure 5.**
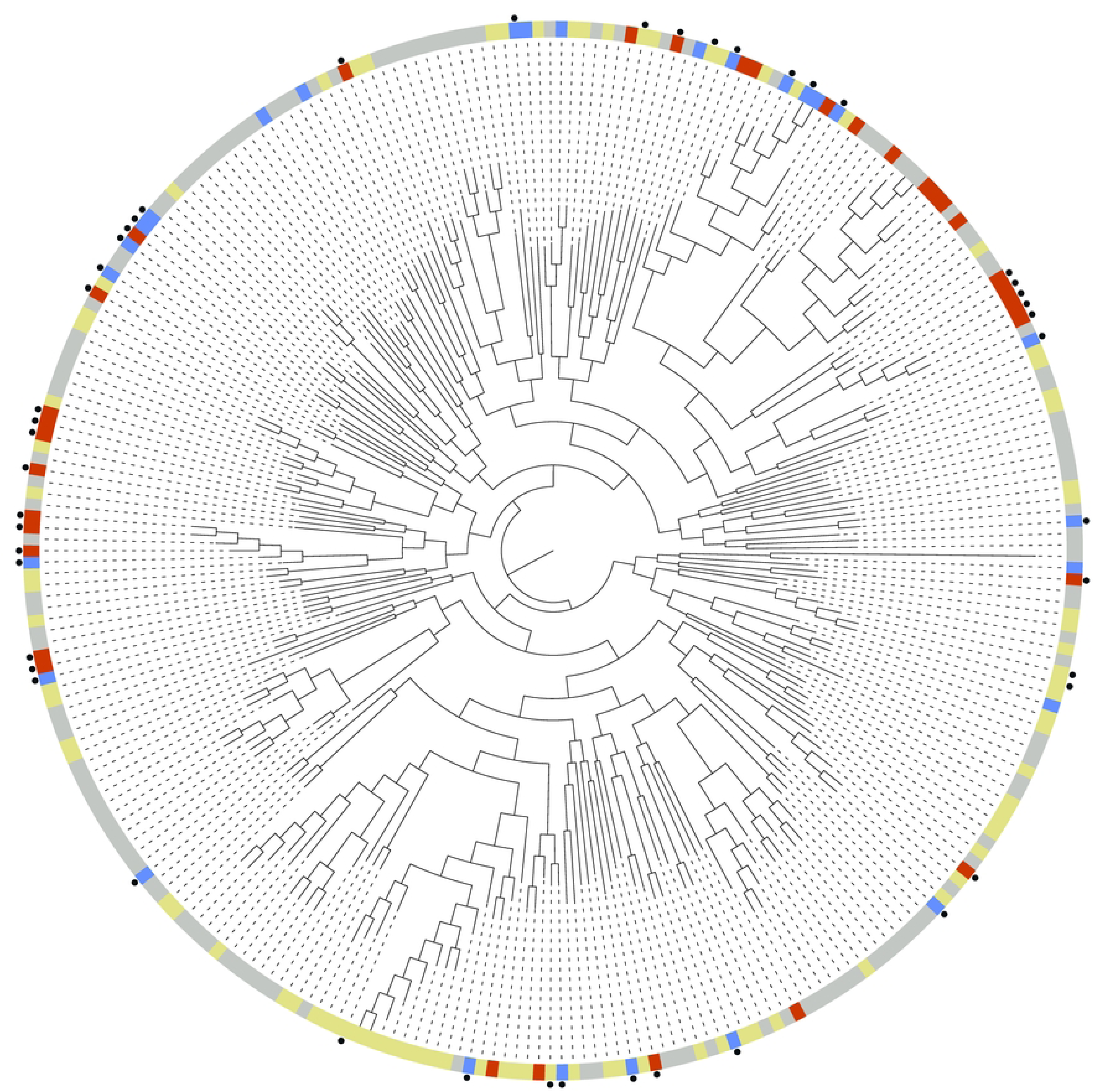
*K. pneumoniae* population structure and phylogenic placement of sequenced *K. pneumoniae* strains. A midpoint rooted maximum-likelihood phylogenetic tree was constructed based on a core gene alignment of 281 *K. pneumoniae* genomes. Isolates sequenced in this study are marked with a black circle. The node color indicates the origin of each isolate as follows: flies (red), manure (blue), mastitis (yellow), other (grey).

### Analysis of sequenced Staphylococcaceae isolates

A total of 145 Staphylococcaceae isolates were sequenced in the study (127 fly-derived, 17 manure-derived, and 1 mastitis-derived), which were identified as belonging to 13 distinct *Staphylococcus* species and 2 distinct *Mammaliicoccus* species. These included isolates from the following species: *Mammaliicoccus lentus* (n= 2), *M. sciuri* (n= 46), *Staphylococcus arlettae* (n= 5)*, S. caeli* (n= 2)*, S. chromogenes* (n= 2)*, S. equorum* (n= 1)*, S. gallinarum* (n= 8)*, S. haemolyticus* (n= 1)*, S. pasteuri* (n= 1)*, S. pseudoxylosus* (n= 8)*, S. saprophyticus* (n= 2)*, S. shinii* (n= 21)*, S. succinus* (n= 3)*, S. ureilyticus* (n= 1), and *S. xylosus* (n= 42) (Table S1).

*M. sciuri* (formally *S. sciuri*) was the most common Staphylococcaceae, with 35 fly-derived strain and 11 manure-derived sequenced. To further analyze the phylogenetic diversity of sequenced *M. sciuri*, all publicly available *M. sciuri* genomes were downloaded from NCBI to create a reference genome set (n=235 after removal of low quality or replicated strains). A phylogenetic tree was constructed based on a core genome alignment of 2,037 orthologs, which showed that most fly-derived strains fell within two clades on the tree (Figure 6A). The tree structure based on the core-genome alignment was compared to a separate outgroup-rooted phylogeny based on sequences of seven housekeeping genes (*ack*, *aroE*, *ftsZ*, *glpK*, *gmk*, *pta1*, *tpiA*) in the *M. sciuri* MLST scheme (Figure S3) (36).

**Figure 6.**
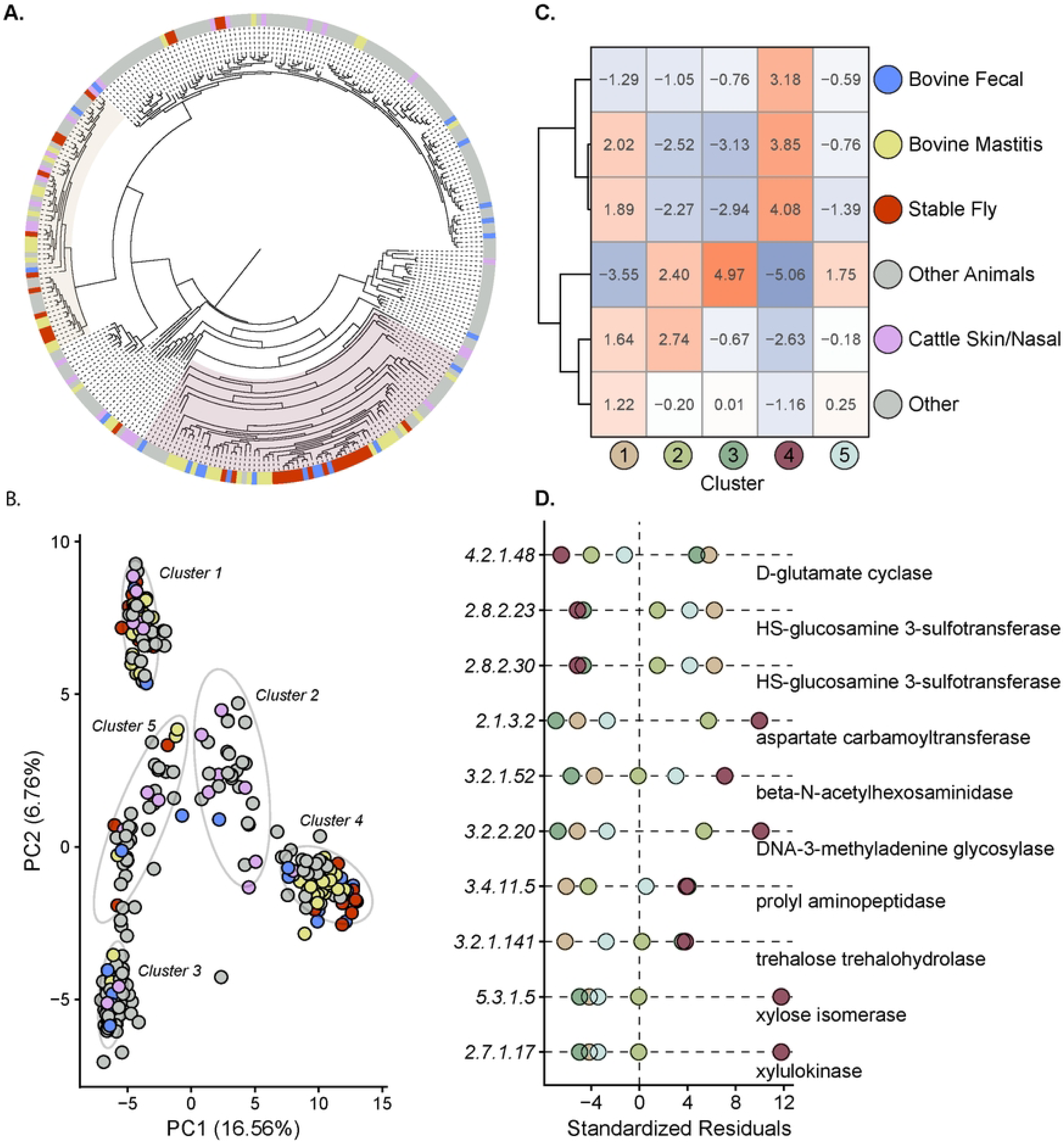
*M. sciuri* genomic analysis. (A) Midpoint rooted maximum-likelihood phylogenetic tree based on core gene alignment of 281 *M. sciuri* genomes, including the 35 fly-derived and 11 manure-derived strains sequenced in this study. Sample origins are colored as follows: flies (red), manure or cattle gastrointestinal tract (blue), bovine mastitis (yellow), cattle skin or nasal (purple), other (grey). The two fly-associated clusters of interest are highlighted respectively on the tree. (B) PCA clustering of genomes based on gene content as determined by Panaroo. Ellipses were calculated with a 95% confidence interval for the five clusters identified through K-means analysis. (C) Heatmap of Chi-Squared standardized residuals (10000 Monte Carlo simulations) showing the association of isolates from different sources with each K-means cluster. The sample origins were further hierarchically clustered based on Euclidean distances, as shown by the dendrogram. (D) Dot-plot of Chi-Squared standardized residuals (10000 Monte Carlo simulations) showing the association of select ECs with *M. sciuri* K-means clusters. Dot-plot shows KEGG ECs for clusters 1 and 4 with a calculated standardized residual value greater than 3.5 and a prevalence greater than 80%. The full list of residual values for all ECs is available in Table S6.

Further analysis revealed that the *M. sciuri* pangenome was largely open with high accessory gene content (Figure S4). We identified that the *M. sciuri* genomes could be grouped into 5 distinct clusters based on gene content using a K-means algorithm (Figure 6B), which largely corresponded to clades on the phylogenetic tree for cluster 1 and cluster 4 (Figure 6A; Figure S3). We then performed a chi-square test with Monte Carlo simulations to determine if there were any significant associations between cluster membership and strain isolation source (overall p-value = 9.999e-05) (Figure 6C). The corresponding standardized residual (R) values indicated that fly-, manure-, and mastitis-derived isolates were most associated with cluster 4 (R > 3), and skin/nasal-derived isolates were more associated with cluster 2 (R=2.74). Bovine mastitis and fly-derived isolates also showed some associations were cluster 1 isolates (R= 2.02 and 1.89, respectively). We used the eggNOG mapper software to perform functional annotation of predicted protein sequences to identify KEGG (Kyoto Encyclopedia of Genes and Genomes) Orthologs (KOs) and Enzyme Commission (ECs) annotations present in each genome (37–39). A chi-square test with Monte Carlo simulations was performed on counts of EC groups, and standardized residuals were used to evaluate the association of each functional group with a specific cluster (overall p-value = 9.999e-05) (Table S6). A total of 10 ECs were found in the fly-associated clusters with a calculated chi-square standardized residual value greater than 3.5 and a prevalence greater than 80% in the respective cluster (Figure 6D). Notably, two ECs related to [heparan sulfate]-glucosamine 3-sulfotransferase were highly associated with cluster 1 but were completely absent from all strains in cluster 4. Conversely, we found that ECs related to Xylose metabolism were highly associated with cluster 4, while being largely absent from genomes assigned to all other clusters (Figure 6D; Table S6).

Construction of individual phylogenies across the Staphylococcaceae family revealed the presence of phylogenetically clustered fly- and manure-derived strains for the following genera: *M. sciuri, S. arlettae*, *S. gallinarum, S. xylosus* (Figure 6A; Figures S5-S9). For each pair, we performed a pairwise genome alignment (MUMmer4) to compare sequence similarity between strains and found several instances of highly similar genome pairs (Table 1, Table S7) (40). We identified three pairs of *S. gallinarum* isolates, which showed on average greater than 99.99% alignment of all bases with a range of 12-351 total unaligned base pairs in the alignment. Of the remaining pairwise alignments (Table 1), the percentage of total aligned base pairs ranged from 95.852% to 99.986%. For the closest *M. sciuri* strain pair (AS0296-AS0565), we identified a total of four contigs unique to the fly-derived strain, which notably included a 3016 bp contig containing a plasmid *rep* and *pre* gene (Table S7). The remaining three contigs were small (<400 bp) with no detected coding sequences (CDS). For the *S. arlettae* strain pair (AS0200-AS0572), the fly-derived strain encoded 4 unique contigs, which notably included a 3,455 bp contig encoding *pre* and *rep* genes as well as a 4,551 bp contig encoding the *pre*, *rep*, and *tet* genes.

**Table 1.** MUMmer4 alignment summary of fly and manure strain pairs clustered together on the core genome phylogenies. Manure strains AS0571 and AS0569 and fly strain AS1286 were obtained from samples collected from the Dairy Cattle Center (DCC), a smaller satellite farm of the main sample collection site. Detailed alignment statistics are available in Table S7.

|  | Fly strain | Bases aligned | Bases unaligned | Manure strain | Bases aligned | Bases unaligned |
| --- | --- | --- | --- | --- | --- | --- |
| *M. sciuri* | AS0296 | 99.7454% | 6815 | AS0565 | 99.9495% | 1350 |
|  | AS1298 | 98.7064% | 33462 | AS0567 | 98.0334% | 51283 |
|  | AS0650 | 95.2483% | 127306 | AS1395 | 98.5790% | 36823 |
|  | AS0913 | 96.5620% | 89859 | AS0579 | 95.8520% | 109361 |
| *S. arlettae* | AS0200 | 99.4203% | 14808 | AS0572 | 99.9797% | 515 |
| *S. gallinarum* | AS0915 | 99.9996% | 12 | AS0571 | 99.9996% | 31 |
|  | AS0915 | 99.9962% | 112 | AS0564 | 99.9951% | 144 |
|  | AS1286 | 99.9880% | 351 | AS0564 | 99.9961% | 113 |
|  | AS1336 | 98.0958% | 56631 | AS0564 | 99.9860% | 408 |
| *S. xylosus* | AS1331 | 98.5428% | 40873 | AS0569 | 98.3096% | 47531 |

### Diverse Staphylococcaceae isolates harbor highly similar ARG encoding plasmid sequences

All sequenced Staphylococcaceae genomes were screened for the presence of both plasmid *rep* genes and ARGs to identify potential plasmid-borne ARGs shared between strains. Of the 19 identified ARGs, 9 genes were found exclusively on contigs without a *rep* gene, indicating either a chromosome origin or that the ARGs are encoded as part of a larger plasmid not identified through short read sequencing. Conversely, *lnuA* and *tetK* were the most common genes encoded on the same contig with a plasmid *rep* and were primarily found as the sole ARG on the contig (Figure 7A; Table S8). Three plasmid replicon proteins (Rep13_7, Rep13_4, Rep21_19) were associated with contigs encoding a single copy of *lnuA* and had a total contig length ranging from 2,126 bp to 5,110 bp (Figure 7B, Figure S10). We performed pairwise-sequence alignments across all 22 *lnuA* encoding contigs and three added reference plasmids via MUMmer4-MobMess (40,41). Contig pairs showed an overall range of 13.27% to 100% global average nucleotide identity; k-means clustering analysis further indicated that contigs were separated into three groups with some intermixing of rep13_7 and rep13_4 plasmids based on overall sequence identity (Figure 7C; Table S8). Notably, a 2,848 bp contig containing *lnu*A and rep13_7 was shared with 100% global average nucleotide identity between seven different strains, which included one fly-derived *M. sciuri* (AS0299), two manure-derived *M. sciuri* (AS0549, AS0554), two fly-derived *S. xylosus* (AS0750, AS1297), one fly-derived *S. arlettae* (AS1300), and one mastitis-derived *S. haemolyticus* (AS0366). This analysis was repeated for plasmid-associated *tetK* contigs, which showed a range of global average nucleotide identity of 76.60% to 100%. We found that 13 of the 16 *tetK* plasmid contigs clustered in one group with two full reference plasmids, which showed a very high degree of sequence similarity (97.30% to 100%; median 99.76%) with similar contig/plasmid lengths (4,439 bp to 4,612 bp) (Figures S11-S12).

**Figure 7.**
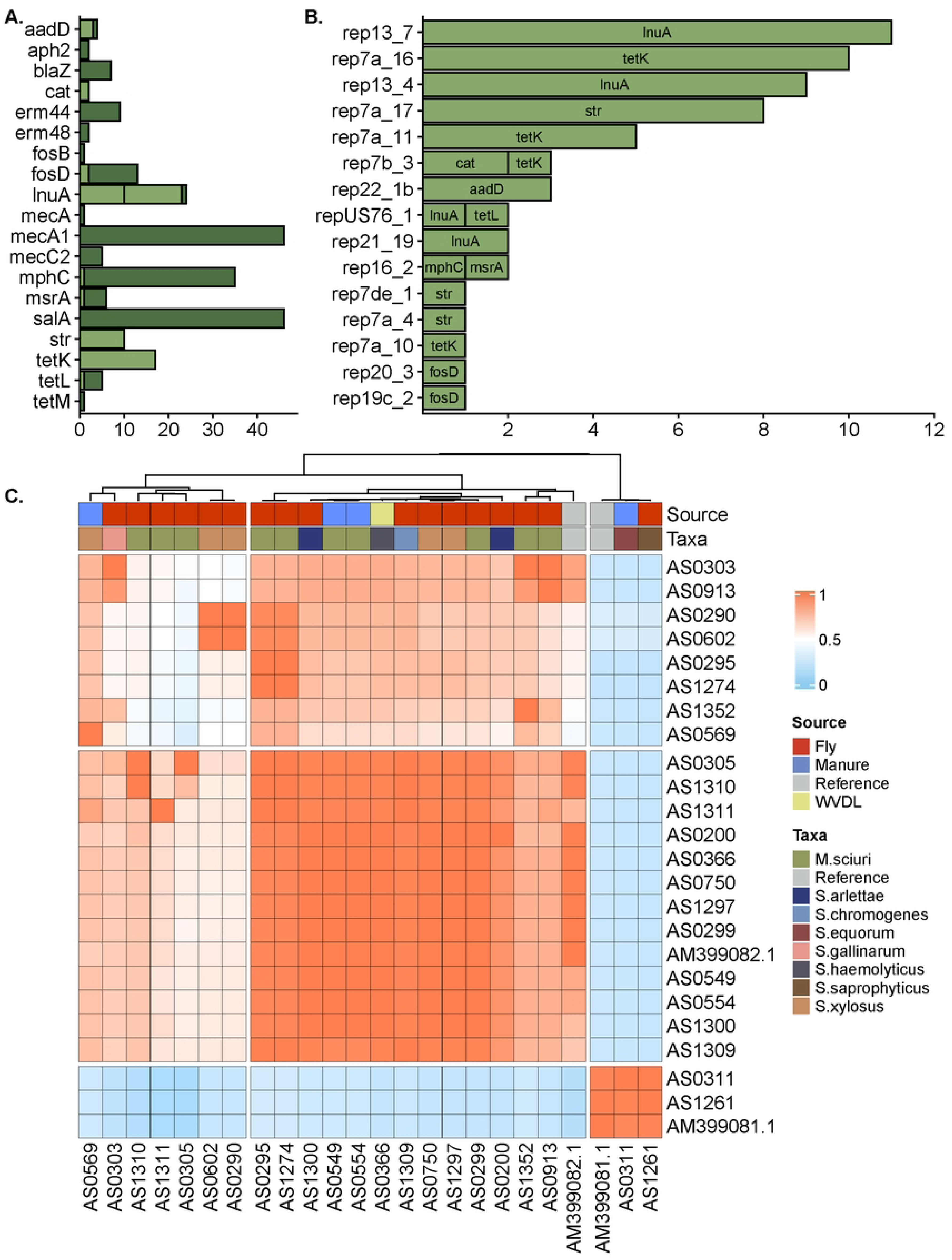
Staphylococcaceae ARG analysis. (A) Bar plot showing the counts of different ARGs found either on the same contig as a plasmid *rep* gene (light green) or on a contig without a *rep* gene (dark green). (B) Bar plot showing the counts of ARGs found on the same contig as different plasmid replicon types, as identified by PlasmidFinder. (C) Heatmap depicting the global percent nucleotide similarity of plasmid-associated contigs encoding the *lnuA* gene. The red indicates a greater percent nucleotide similarity while blue indicates a lower percent similarity. Associated strain name is labelled for both the row and axis, with AM399082.1 and AM399081.1 included as reference plasmids. Both rows and columns were hierarchically clustered based on Euclidean distance, with only the column dendrogram shown. The origin and taxonomic identification of the strains are marked below the dendrogram. Four strains (AS0311, AS0554, AS0569, AS1261) were isolated from samples collected from the Dairy Cattle Center (DCC), a smaller satellite farm of the main sample collection site.

## Discussion

The overarching goal of this study was to examine the potential ecological flow of manure-associated bacterial isolates to hematophagous *Stomoxys* flies. *Stomoxys* populations on livestock facilities are sustained by a near constant availability of both mammalian hosts and manure, which serve as important developmental and nutritional resource for biting muscid flies (12–16). Manure also serves as a major reservoir human and bovine pathogens, including opportunistic Enterobacteriaceae and Staphylococcaceae bacteria responsible for udder tissue inflammation (bovine mastitis) in lactating cattle (1). Both culture based and amplicon sequencing based studies have demonstrated that the *Stomoxys* microbiota is largely composed of opportunistic pathogens (7–10); however, genomic based studies are still needed to validate the connection between biting flies and manure reservoirs.

A key aim of this study was to determine the genomic diversity of fly-derived bacteria in relation to strains associated with cattle hosts. For *E. coli*, we observed that fly- and manure-derived isolates were interspersed along the reconstructed phylogenetic tree, suggesting the absence of large scale *Stomoxys*-exclusive *E. coli* lineages in the sequenced samples. Instead, we found that the majority of both fly- and manure-derived *E. coli* strains were assigned to phylogroup B1 or A, which are often associated with a commensal niche in the bovine digestive tract (42–45). Notably, we did detect a minority of strains encoding Afa/Dr adhesion genes, which are the primary PCR diagnostic targets for the diffusely adherent *E. coli* (DAEC) diarrheagenic pathotype (46). A subset of fly strains also possessed the F17 family fimbriae, which facilitate bacterial attachment to the bovine intestinal mucosal cells and have been previously linked to enteric and septicemic infections in calves (47,48). These strains, however, do not encode any identified heat-labile toxins (LT), which are also key virulent determinants of enterotoxigenic *E. coli* (ETEC) (49). Nevertheless, a subset of phylogroup A and B1 strains encoded the high pathogenicity island (HPI), a biosynthetic gene cluster responsible for the production of the yersiniabactin siderophore, a high virulence determinant originally characterized in *Yersinia enterocolitica* (50,51).

The single mastitis-derived *E. coli* in the sequenced collection was shared the same ST as both a manure and a fly-derived strain, and notably had few identified VFs. These findings support previous work suggesting that mastitis-causing *E. coli* lack specific VFs and instead represent opportunistic environmental strains (52,53). Comparison of our isolates to a larger reference phylogeny showed that the majority of published mastitis-associated *E. coli* strains were assigned to phylogroups A or B1, further supporting the link between bovine mastitis and bovine gastrointestinal tract *E. coli*. We additionally sought to compare our sequenced *E. coli* strains to previously published isolates derived from other fly taxa. To date, several studies have used whole-genome sequencing to analyze *E. coli* carried by nonbiting fly populations, including *Musca domestica* (Family: Muscidae) (54,55), *Chrysomya* sp. (Family: Calliphoridae) (56), or unidentified fly taxa (57–61); however, these samples were collected from non-dairy farm environments (swine farms, poultry farms, or hospitals). The inclusion of these isolates in our reconstructed phylogenetic tree revealed significant diversity in fly-derived *E. coli*, which were likely acquired by individual flies through interactions in the respective environments.

Sequenced *K. pneumoniae* isolates formed three distinct phylogenetic clades, with intermixing of fly-, manure-, and mastitis-derived isolates. Apart from capsule type, there were few differences in identified virulence factors between strains, including for mastitis-derived isolates. Both the capsular polysaccharide (CPS) and the Lipopolysaccharide (LPS) O antigen are important virulence determinants in *K. pneumoniae*, playing key roles in mucosal colonization and evasion of host immune responses (62). We identified a manure-derived strain encoding the K1 capsule type, which is a predominant capsule type found in many human pathogenic *K. pneumoniae* (63). While the polysaccharide capsule is also thought to be important to facilitating invasion of bovine mammary glands, studies often report a great diversity of CPS and LPS encoded by mastitis-derived strains (64,65). Our species-wide phylogenomic analysis also revealed that both fly, manure, and mastitis strains were intermixed with human-derived isolates. The fly-associated strains were not restricted to a single subclade, indicating that muscid flies can be colonized by diverse isolates of *K. pneumoniae*. Our pangenome analysis showed no large-scale clustering of bacterial isolates by gene content, which could indicate a large accessory genome and frequent horizontal transfer between divergent *K. pneumoniae* strains. Alternatively, our analysis may be limited by the lack of genomes available from environmental or non-human sources, which represent a small subset of all sequenced *K. pneumoniae* strains (66).

A major limitation hindering further analysis of many non-*aureus* Staphylococcaceae species is the lack of publicly available genomes for sequence comparison. Despite their role as the most frequent causative agents of subclinical bovine mastitis (23), little is known about the genomic and functional diversity of coagulase-negative Staphylococcaceae species. Here, we performed a comparative analysis of *M. sciuri* (formally *S. sciuri*), the most common *Staphylococcaceae* species sequenced in this study. The inclusion of all publicly available *M. sciuri* genomes revealed strong co-clustering of fly, manure, and mastitis strains based on gene content. This manure-associated cluster also corresponded to a specific clade in the species-wide phylogeny based on a core genome alignment. While xylan hydrolase (*xynB*) was found across nearly all *M. sciuri*, genomes within the manure-associated cluster showed significant enrichment for metabolic genes responsible for the conversion of D-xylose into D-Xylulose (*xylA*) and D-Xylulose 5-phosphate (*xlyB*), which were found to be rare or absent across strains from all other *M. sciuri* clusters. Xylose is a key component of hemicellulose, a major plant fiber component found in the feces of ruminants (67–69), and the ability to metabolize xylose via other pathways could provide a competitive advantage in the bovine gastrointestinal tract (fecal) niche. These results provide further evidence for the role of manure-associated strains as causative agents of mastitis and as prominent members of the *Stomoxys* microbiome. While these findings provide novel insights into potential niche differentiation in *M. sciuri*, further work, including large scale parallel sampling of skin and manure microbes, will be needed to validate these hypotheses.

In this study, we sequenced strains primarily originating from a large free stall barn system; however, a subset of isolates was obtained from a nearby smaller tie-stall barn. While these barns are geographically separated, cattle are frequently moved between the two facilities, and are part of an operationally connected herd management system (7). Pairwise genome comparisons of phylogenetically clustered isolates revealed near identical strains (Up to 99.996% base-pair alignment) derived from flies and manure. This was especially notable for a trio of *S. gallinarum* isolates, including one obtained from a manure sample from the satellite facility, which showed, on average, fewer than 100 base pairs unaligned across the entire draft genomes. We also observed instances where the respective alignments differed primarily by whole contigs containing *rep* genes or other plasmid-associated genes, indicating that sequence divergence among some strains was driven by mobile genetic elements. Collectively, these results suggest that strains of *Staphylococcus* bacteria and their associated mobile genetic elements can be circulated through the dairy barn environment by *Stomoxys* flies, with the potential for cross-facility dissemination. A second major goal of this study was therefore to determine the potential dispersal of plasmid-borne ARGs through the farm environment. In *Enterobacteriaceae*, ESBLs, which are often encoded on plasmids or other mobile genetic elements, provide resistance to clinically significant beta-lactam antibiotics used in human and veterinary medicine (70). We detected ESBLs (*bla*-TEM, *bla*-CTX, *bla*-CMY) in sequenced *Enterobacter*, *Klebsiella*, and *E. coli* genomes; however, only four ESBLs were found on the same contig as a plasmid replicon. We previously reported that these same strains exhibited cephalosporin resistance, which could be transferred to a naïve strain via conjugative transfer *in vivo* (24). The current results therefore likely represent a limitation of short-read sequence data, which is not suitable for the direct assembly of large or complex plasmid sequences (71). In contrast, *Staphylococcus* are known to carry ARGs on small plasmids, which are more effectively captured by short-read sequencing (71–73). The *lnuA* and *tetK* genes were the most common ARGs encoded on contigs with a plasmid *rep* gene, which showed strong structural similarity with known reference plasmid sequences. Analysis of these sequences showed that near-identical plasmids were shared between fly and manure isolates and across multiple taxa, further providing evidence for the dispersal of ARG plasmids across the environment.

Collectively, our results provide evidence for the circulation of both individual bacterial strains and plasmids between flies and bovine manure on dairy farms. This study, while likely an underrepresentation of the total strain diversity in the environment, still represents the first comparative genomic analysis of *Stomoxys*-derived bacterial isolates. Our study provides strong evidence for the acquisition of fecal bacteria by muscid flies on dairy farms, and suggests that the microbiomes of flies and manure are highly interconnected. Given the role of manure as a key reservoir of both mastitis and zoonotic pathogens(1–3), biting flies may represent important mechanical vectors of manure pathogens and could pose a biosafety hazard on dairy farms.

## Materials and Methods

### Collection and sequencing of bacterial isolates

Bacterial isolates sequenced in this study were collected as part of three previous studies to characterize the microbiota of biting *Stomoxys* flies in relation to environmental manure piles (7,8,24). Detailed methodology related to the collection of *Stomoxys* flies and environmental manure as well as subsequent bacterial isolations are reported in these studies. Here, we selected a subset of these isolates for short-read sequencing, with representative isolates selected from each unique fly pool and manure sample. DNA from the selected isolates was extracted using a NucleoSpin Tissue kit (Macherey-Nagel, Nordrhein-Westfalen, Germany), and quantified using with a Quantus fluorometer (Promega, Madison, WI, USA). Genomic DNA sent to the SeqCoast Genomics (Portsmouth, NH, USA) were prepared using the Illumina DNA Prep tagmentation kit (San Diego, CA, USA) and sequenced on the NextSeq2000 platform using a 300 cycle flow cell kit to produce 2x150bp paired reads with an estimated 1.3 million reads per sample. Read demultiplexing and read trimming were performed using DRAGEN v3.10.12 as part of the SeqCoast Genomics pipeline. Genomic DNA sent to the UW-Madison Biotechnology Center were prepared using the Celero EZ DNA-Seq prep kit (Tecan Genomics, Männedorf, Switzerland) and sequenced on the Illumina NovaSeq6000 platform to generate 2x150bp shared reads with an estimated 1 million reads per sample. Removal of adapter and low-quality sequences was performed using via Trimmomatic (v0.39) (74). Sequence quality was checked on both raw and trimmed reads using FastQC (v0.11.9) (75).

### Genome assembly, annotation, and quality control

Genome assembly was performed using the SPAdes assembler (v3.15.5) (76) as implemented through Shovill (v1.1.0) (77). Assembly statistics were determined using QUAST (v5.2.0) and detection of expected universal single copy orthologs was performed using BUSCO (v 5.7.1) (25,26). The completeness and contamination levels of draft assemblies were assessed using the CheckM (v1.0.18) full reference tree lineage workflow on the KBase platform (78,79). All assembly statistics are available in the supplemental files (Table S1). Contigs were annotated using the Prokka pipeline (v 1.13) using the optional ‘rfam’ computation setting (80).

### Taxonomic classification and typing of isolates

Identification of bacterial isolates were performed through comparison of draft assemblies to the PubMLST Ribosomal Multilocus Sequence Typing (MLST) database, which resolves taxonomy based on 53 ribosome protein subunits (*rps*) genes (81). For identified *E. coli* isolates, sequence types were determined through comparison of allelic profiles against the Pasteur eight gene MLST scheme and the phylogroup was predicted using the ClermonTyping program (32,82). Further *in silico E. coli* serotyping was performed via the Center for Genomic Epidemiology (CFGE) SerotypeFinder (v 2.0.1) (83). For isolates identified as *K. pneumoniae,* Kleborate (v2.4.1; Kaptive setting) was used to determine the sequence type and to predict the K (capsule) and O antigen (LPS) serotype (35,84). ABRicate (v1.01; databases downloaded 2024-Jun-11) was used to screen all contigs for the presence of plasmid replicon genes (*rep*) (PlasmidFinder), VFs (VFDB; VirulenceFinder for *E.coli* isolates), and ARGs (Resfinder) (27–31).

### Phylogenetic analysis in a local and global context

Phylogenetic analysis was performed for the most highly represented taxonomic groups in the collection (*E. coli, K. pneumoniae*, *M. sciuri*, and *Staphylococcus spp.*). Panaroo (v1.5.0) was used to produce a core genome alignment (core_threshold 0.98; MAFFT) files for each listed taxonomic group (85,86). Maximum-likelihood phylogenetic trees were inferred on filtered core genome alignments using a Jukes-Cantor + CAT nucleotide substitution model via FastTree (v2.1.11) (87). Reference genome sets were constructed for each taxonomic group to determine the global phylogenetic placement of sequenced isolates (Table S2). For *E. coli*, a reference genome set was constructed based on available genomes from the *Escherichia coli* reference collection (ECOR) (88,89), and further supplemented with select pathotype genomes as well as mastitis- and fly-derived strains. For *K. pneumoniae*, genomes were compiled from a previously described reference set along with available mastitis- and fly-derived strains (66). For both *M. sciuri* and *Staphylococcus spp.*, the initial reference sets included all respective publicly available genomes from the NCBI Genome database. The drep genome comparison tool (dRep compare --P_ani .999 --S_ani .999; v3.5.0) was used to identify highly similar genomes within each reference set based on both MASH and ANI (90). To minimize duplication of strains, a representative strain was selected when two or more genomes from the same environment and submission details (BioProject) shared more than 99.99% average nucleotide identity as calculated through fastANI (v 1.33) (91). Global phylogenies were constructed based on annotated genomes using Panaroo and FastTree, as described above (80,85,87). Bacterial phylogenies were visualized in the ‘ggtree’ package and the TreeViewer software.

### Identification of clustering based on pangenome features and functional annotations

Bacterial pangenome features were imported into R for further analysis using the ‘Pagoo’ package (85,92). A principal component analysis was performed on the pan genome matrix using the ‘pagoo::pan_pca()’ function and associated clustering analysis was performed using k-means algorithm via the ‘factoextra’ package (v1.0.7) (93). To test for host association with specific clusters, we performed chi-square tests (simulated p-values with 10000 Monte Carlo simulations) and visualized the standardized residuals via the ‘pheatmap’ (v1.0.12) package in R (94).

Further functional annotation was performed to identify orthologs associated with identified bacterial clusters or lineages. Predicted protein sequences were used as an input to eggNOG-mapper (v2.1.12) to perform a diamond BLAST (v2.1.10) sequence alignment against the eggnog protein database (v5.0.2) with a 0.001 minimum hit e-value (37–39). Resulting annotation files were imported into R to compute counts of identified KEGG Enzyme (EC) numbers (95,96). Chi-square tests (simulated p-values with 10000 Monte Carlo simulations) were performed on calculated contingency tables to test for significant association between functional counts and taxonomic clusters. Descriptive information for significant ECs were attached using the ‘KEGGREST’ package and calculated standardized residuals were visualized using ‘ggplot2’(97,98).

### Identification of highly similar Staphylococcaceae strains and plasmids

We further screened phylogenetic trees to identify clustered pairs of phylogenetically related sequenced isolates from different sources (fly, manure, mastitis). To determine strain similarity, pairwise genome comparisons and were performed on selected pairs using Mummer4(nucmer v4.0.0, DNAdiff v1.3) (40). To determine the potential transmission of small ARG-encoding plasmids throughout the environment, contig sequences containing both a plasmid replicon gene and an associated ARG were extracted from *Staphylococcaceae* draft genome assemblies. For comparison, contigs were re-oriented to include all *rep* genes in the forward direction. Plasmids were sorted by *rep* type (as identified above by PlasmidFinder) and subsequent pairwise sequence alignments to determine percent similarity were performed using MUMmer4 (v4.0.0) as implemented MobMess (29,40,41). The resulting pairwise global percent similarities were visualized in R using ‘complex heatmap’ (99). Plasmid maps were constructed using ‘gggenomes’ on reindexed annotated (Bakta v1.8.2) contig sequences to include *rep* genes in the starting position for easier visualization (100,101).

### Data availability

Raw Illumina reads and assemblies are available in the NCBI Sequence Read Archive (https://www.ncbi.nlm.nih.gov/sra) under BioProject ID PRJNA1216036. Data supporting the conclusions of this article, along with scripts used for analysis and figure generation, are provided as supplemental materials or are available in the Coon laboratory’s GitHub repository (https://github.com/kcoonlab/stable-fly-genomes). Physical isolates are available upon request.

Reviewer link to NCBI data: https://dataview.ncbi.nlm.nih.gov/object/PRJNA1216036?reviewer=h3engda705p6us7kcdrb2hdgq8

## Acknowledgements

We thank Garret Suen (Department of Bacteriology, University of Wisconsin-Madison) for advice regarding framing and presentation of this work. We also thank Jessica Cederquist and the UW-Madison Department of Animal and Dairy Science Dairy Herd Operation for access to the dairy cattle facilities included in this study (Arlington and the DCC), along with their associated records, as well as the Wisconsin Veterinary Diagnostic Laboratory for sharing select mastitis strains used in this study.

## Supporting information

**Figure S1.** *E. coli* pangenome analysis. (A) Pan and core genome curves as calculated by Pagoo. (B/C) PCA visualizations based on gene content. Figure S1B is colored by sample origin as follows: flies (red), manure (blue), mastitis (yellow), other (grey). The dark red dots indicate isolates sequenced in this study; the light red dots correspond to publicly available fly-derived *E. coli* genomes. Figure S1C is colored by MASH phylogroup as follows: D (Dark blue), B2 (cyan), E (orange), A (light red), B1 (green), other (grey).

**Figure S2.** *K. pneumoniae* pangenome analysis. (A) Pan and core genome curves as calculated by Pagoo. (B) PCA visualizations based on gene content. The color corresponds to the sample origin as follows: flies (red), manure (blue), mastitis (yellow), human (dark grey), other (grey). The dark red dots indicate isolates sequenced in this study; the light red dots correspond to publicly available fly-derived genomes.

**Figure S3.** *M. sciuri* supporting phylogenetic trees. (A) Outgroup rooted maximum-likelihood phylogenetic tree based on alignment of six housekeeping genes. Tree is rooted with an *S. xylosus* isolate as an outgroup. (B) Midpoint rooted maximum-likelihood phylogenetic tree based on alignment of core genes. For both trees, the exterior rectangle node annotation corresponds to the sample origin and the interior node annotation corresponds to the cluster as assigned by gene content (See Figure 6 legend).

**Figure S4**. *M. sciuri* pangenome analysis supporting figures. (A) Pan and core genome curves as calculated by Pagoo. (B) Barplot showing the frequency distribution of identified gene clusters in the *M. sciuri* pangenome.

**Figures S5-S9.** Phylogenetic trees of other Staphylococcaceae isolates. All phylogenies are shown as midpoint rooted maximum-likelihood trees based on an alignment of core genes within each genus.

**Figure S10.** The genomic architecture of *lnuA* encoding plasmids. The figure depicts schematic diagrams indicating the location of *lnuA*, *rep* (replicon protein gene), and *tnp* genes on plasmids identified from sequenced *Staphylococcaceae* strains along with selected reference sequences. All sequences were reindexed to position the rep gene on the left-hand side of the forward strand. The rep gene variant (identified via PlasmidFinder) is noted.

**Figure S11.** Heatmap depicting the global percent nucleotide similarity of plasmid-associated contigs encoding the *tetK* gene. Percent similarity and associated strain metadata (taxonomic identification, strain origin) are marked as described in Figure 7. MH785224.1, CP094730.1, and CP038249.1 are included as reference strains.

**Figure S12.** The genomic architecture of *tetK* encoding plasmids. The figure depicts schematic diagrams indicating the location of *tetK*, *rep* (replicon protein gene), and *pre* (plasmid recombination enzyme) genes on plasmids identified from sequenced Staphylococcaceae strains along with selected reference sequences. All sequences were reindexed to position the rep gene on the left-hand side of the forward strand. The *rep* gene variant (identified via PlasmidFinder) is noted on the right-hand side.

**Table S1.** Genome assembly and quality statistics

**Table S2.** Screening of all contigs for VFs, ARGs, and plasmid *rep* genes

**Table S3.** Further typing of *E. coli* isolates, and associated metadata related to Figure 3

**Table S4.** Construction of reference sets for phylogenetic placement of isolates

**Table S5.** Further typing of *K. pneumoniae* isolates, and associated metadata related to Figure 5

**Table S6.** Chi-Square residuals and prevalence data for *M. sciuri* KEGG ECs

**Table S7.** MUMmer4 pairwise genome analysis and analysis of unaligned contigs

**Table S8.** Co-occurrence of ARGs and plasmid *rep* genes in *Staphylococcaceae* isolates

## Author contributions

### Conceptualization

Ideas; formulation or evolution of overarching research goals and aims. *AJS, KLC*

### Data Curation

Management activities to annotate (produce metadata), scrub data and maintain research data (including software code, where it is necessary for interpreting the data itself) for initial use and later reuse. *AJS, KLC*

### Formal Analysis

Application of statistical, mathematical, computational, or other formal techniques to analyze or synthesize study data. *AJS*

### Funding Acquisition

Acquisition of the financial support for the project leading to this publication. *AJS, KLC*

### Investigation

Conducting a research and investigation process, specifically performing the experiments, or data/evidence collection. *AJS*

### Methodology

Development or design of methodology; creation of models *AJS, TWK, PS, KLC*

### Project Administration

Management and coordination responsibility for the research activity planning and execution. *KLC*

### Resources

Provision of study materials, reagents, materials, patients, laboratory samples, animals, instrumentation, computing resources, or other analysis tools. *KLC*

### Software

Programming, software development; designing computer programs; implementation of the computer code and supporting algorithms; testing of existing code components. *AJS*

### Supervision

Oversight and leadership responsibility for the research activity planning and execution, including mentorship external to the core team. *KLC*

### Validation

Verification, whether as a part of the activity or separate, of the overall replication/reproducibility of results/experiments and other research outputs. *AJS, KLC* **Visualization** Preparation, creation and/or presentation of the published work, specifically visualization/data presentation. *AJS*

### Writing – Original Draft Preparation

Creation and/or presentation of the published work, specifically writing the initial draft (including substantive translation). *AJS*

### Writing – Review & Editing

Preparation, creation and/or presentation of the published work by those from the original research group, specifically critical review, commentary or revision – including pre- or post-publication stages. *AJS, TKW, PS, KLC*

*All authors read and approved the final version of the manuscript*.

